# Development of a highly-sensitive method to detect the carriage of carbapenem-resistant *Pseudomonas aeruginosa* in humans

**DOI:** 10.1101/2024.08.27.609846

**Authors:** Selvi N. Shahab, Anneloes van Veen, Nikita Kempenaars, Amber Rijfkogel, Heike Schmitt, Yulia R. Saharman, Margreet C. Vos, Anis Karuniawati, Juliëtte A. Severin, the SAMPAN Consortium

## Abstract

Carbapenem-resistant *Pseudomonas aeruginosa* (CRPA) causes severe and potentially life-threatening infections in hospitalized patients with mortality rates of more than 40%. To detect CRPA carriage in humans for surveillance purposes or to prevent spread and outbreaks in hospitals, a highly-sensitive culture method for CRPA carriage in humans is needed. We aimed to develop such a highly-sensitive method, that would be feasible in laboratories with limited resources. In this study, seven well-defined CRPA strains belonging to high-risk clones were used, including one CRPA without a carbapenemase gene and six carbapenem-resistant isolates with carbapenemase genes. We applied a stepwise approach wherein we included four enrichment broths and eight *Pseudomonas aeruginosa*-selective culture media. Spiking experiments were performed to further evaluate the combination of the most sensitive enrichment broths and selective agar plates in human samples. The two most sensitive enrichments broths were TSB-vancomycin and TSB-vancomycin with 2 mg/L imipenem and the most sensitive selective agar plates were *Pseudomonas* isolation agar Becton Dickinson, *Pseudomonas* isolation agar Sigma-Aldrich, and M-PA-C (Becton Dickinson). After the spiking experiment, the best method for detecting CRPA based on the sensitivity and the selectivity was the combination of TSB-vancomycin with 2 mg/L imipenem as an enrichment broth for overnight incubation, followed by subculturing the broth on M-PA-C agar plate. We have thus developed a highly-sensitive selective method to detect CRPA carriage in humans, which can also be applied in limited-resource laboratories. This may contribute to an overall effort to control CRPA.

## INTRODUCTION

*Pseudomonas aeruginosa* is a Gram-negative bacterium that causes severe and potentially life-threatening infections in hospitalized patients (1). The worldwide emergence of carbapenem-resistant *P. aeruginosa* (CRPA) makes infections by these pathogens almost untreatable, resulting in crude mortality rates of more than 40% (2, 3). The World Health Organization has, consequently, ranked CRPA as “high priority” bacterial pathogen for further action (4).

In the hospital setting, actions should be focused on the prevention of transmission of CRPA. Several studies have reported colonization with CRPA in admitted patients, which poses a risk of transmitting these pathogens to other patients or environmental reservoirs where these bacteria may be difficult to eradicate and lead to outbreaks (2, 5, 6). During outbreaks, contact investigations should be performed to identify undetected carriers (7). For these purposes, a highly-sensitive culture method for CRPA carriage in humans is needed. Retrieving viable isolates is essential for antimicrobial susceptibility testing and, if available, analysis of genetic relatedness among isolates (8, 9).

In a recent review on this topic (10), a lack of knowledge on the methods to be used for the rapid and sensitive detection of CRPA was revealed, which was reflected by only a few diagnostic accuracy studies comparing different culture methods and a large variety of culture methods described in recent outbreak-surveillance studies. It was suggested, however, that the use of an enrichment broth prior to plating the material on a selective medium would be of benefit, although this was based on only one study. Therefore, the aim of this study was to develop a highly-sensitive culture method for the detection of CRPA carriage in humans. To that end, various enrichment broths (i.e., tryptic soy broth [TSB] with addition of various antibiotics) and *P. aeruginosa-*selective agar plates were compared, followed by spiking experiments to determine the most sensitive combination of broth and plate to detect CRPA. We aimed to develop a method that would be feasible in laboratories with limited resources as well, thus without application of a nucleic acid amplification technology.

## MATERIALS AND METHODS

### General approach

The study was performed in the framework of the SAMPAN study (A Smart Surveillance Strategy for Carbapenem-resistant *Pseudomonas aeruginosa*) (11). We developed the method in a stepwise manner, by

- Testing the growth of well-characterized CRPA strains inoculated into an enrichment broth continued by culturing onto blood agar
- Testing the phenotype and growth of well-characterized CRPA strains on various *P. aeruginosa* selective agar plates
- Evaluating the growth of well-characterized CRPA strains from faecal samples spiked with these strains while using the most sensitive enrichment broth from previous experiments continued by culturing onto the most sensitive selective agar plates

### Bacterial isolates

In this study, seven well-defined CRPA strains from Indonesia and the Netherlands were used, including one CRPA without a carbapenemase gene and six carbapenem-resistant isolates with carbapenemase genes, including *bla*_VIM_ (n=3), *bla*_GES_ (n=1), *bla*_IMP_ (n=1), and *bla*_NDM_ (n=1) (Table 1). Species identification was performed using the Matrix-Assisted Laser Desorption/Ionization Time-Of-Flight mass spectrometry (MALDI-TOF MS) (Bruker Daltonics, Bremen, Germany). Antibiotic susceptibility was determined by VITEK2® (bioMérieux, Marcy l’Etoile, France) (12). Carbapenem-resistance was defined as resistance to at least one of the carbapenems (*i*.*e*., imipenem, or meropenem). The results of the susceptibility test were interpreted according to the breakpoints defined by the European Committee on Antimicrobial Susceptibility Testing (EUCAST) (13, 14). Multiplex real-time PCR was performed to detect resistance genes, followed by sequencing to genetically characterize the strains. To make series of bacterial suspensions, inoculums of 0.5 McFarland standard suspensions (approximately 1.5 × 10^8^ CFU/mL) from each of the CRPA strains were prepared using 0.45% saline. Eight times 10-fold serial dilution from each inoculum suspension was made in a 0.45% sterile saline solution creating bacterial suspensions with different concentration from 1.5 × 10^8^CFU/mL until 1.5 CFU/mL. To confirm the purity of the suspensions, 100 µL of the suspensions were inoculated onto tryptic soy agar with 5% sheep blood (i.e., blood agar) (Becton Dickinson Diagnostics, Breda, The Netherlands), followed by spreading the inoculum evenly over the surface of the plate using sterile disposable spatula. The plates were observed the next day to ensure purity and for counting.

**Table 1.** Carbapenem-resistant *Pseudomonas aeruginosa* strains used in this study.

|  | Sequence type | Carbapenem-resistant gene | Minimum Inhibitory Concentration (mg/L) |  |  |
| --- | --- | --- | --- | --- | --- |
|  |  |  | Imipenem | Meropenem | Ceftazidime |
| Strain 1<sup>1</sup> | ST446 | *bla*<sub>VIM-2</sub> | $\geq 16$ (R) | 16 (R) | 16 (R) |
| Strain 2<sup>2</sup> | ST773 | *bla*<sub>NDM</sub> | $> 32$ (R) | $> 32$ (R) | $> 16$ (R) |
| Strain 3<sup>3</sup> | ST111 | *bla*<sub>VIM-2</sub> | $> 8$ (R) | 4 (I) | $\geq 32$ (R) |
| Strain 4<sup>4</sup> | ST253 | *bla*<sub>VIM-2</sub> | $\geq 16$ (R) | $\geq 16$ (R) | $\geq 16$ (R) |
| Strain 5<sup>5</sup> | ST357 | *bla*<sub>IMP-7</sub> | $\geq 16$ (R) | $\geq 16$ (R) | $\geq 32$ (R) |
| Strain 6<sup>5</sup> | ST235 | *bla*<sub>GES-5</sub> | $\geq 16$ (R) | $\geq 16$ (R) | 16 (R) |
| Strain 7<sup>5</sup> | ST446 | None | $\geq 16$ (R) | $\geq 16$ (R) | 8 (S) |

<sup>1</sup>Previously published by Van der Zee et al. (15)
<sup>2</sup>From a patient that was hospitalized in Morocco, and was screened after being transferred to the Netherlands.
<sup>3</sup>Previously published by Pirzadian et al. (16)
<sup>4</sup>Strain from a sink drain in the intensive care, previously published by Pirzadian et al. (17)
<sup>5</sup>Previously published by Pelegriin et al. (18)

### Comparison of enrichment broths

Four enrichments broths, all based on tryptic soy broth (TSB) (Becton Dickinson Diagnostics, Breda, The Netherlands) were included in this comparison (Figure 1): TSB supplemented with 2 mg/L of vancomycin (*i*.*e*., TSB-vancomycin), TSB-vancomycin supplemented with 2 mg/L and 4 mg/L of imipenem, and TSB-vancomycin with 6 mg/L of ceftazidime. Vancomycin was added in all broths, as this inhibits Gram-positive bacteria that are present in perianal, rectal swabs or faeces, which are the most used screening samples (6, 10). As a carbapenem antibiotic, imipenem was used because imipenem is more stable than meropenem when using discs to prepare the broth (19). Ceftazidime was chosen based on the general finding that most CRPA are also less susceptible to ceftazidime, and previous experiences (6). The enrichment broths were prepared by adding antibiotic discs to the broth. For instance, to attain 4 mg/L of vancomycin, two discs of 5 µg (Oxoid) were added to 5mL broth. To compare the enrichment broths, 100 µL of the bacterial suspension dilutions were inoculated into the broths. Subsequently, each broth was incubated at 35±1°C for 24 hours, followed by observation of turbidity. After that, regardless of the turbidity, 10 µL of the broth was sub-cultured onto a blood agar plate, which was incubated at 35±1°C for 24 and 48 hours. Blood agar plates were observed for bacterial growth and each plate was scored as either positive (growth) or negative (no growth). All experiments were performed in triplicate. The sensitivity was calculated as the number of positive plates divided by the total number of samples tested per type of broth.

**Figure 1.**
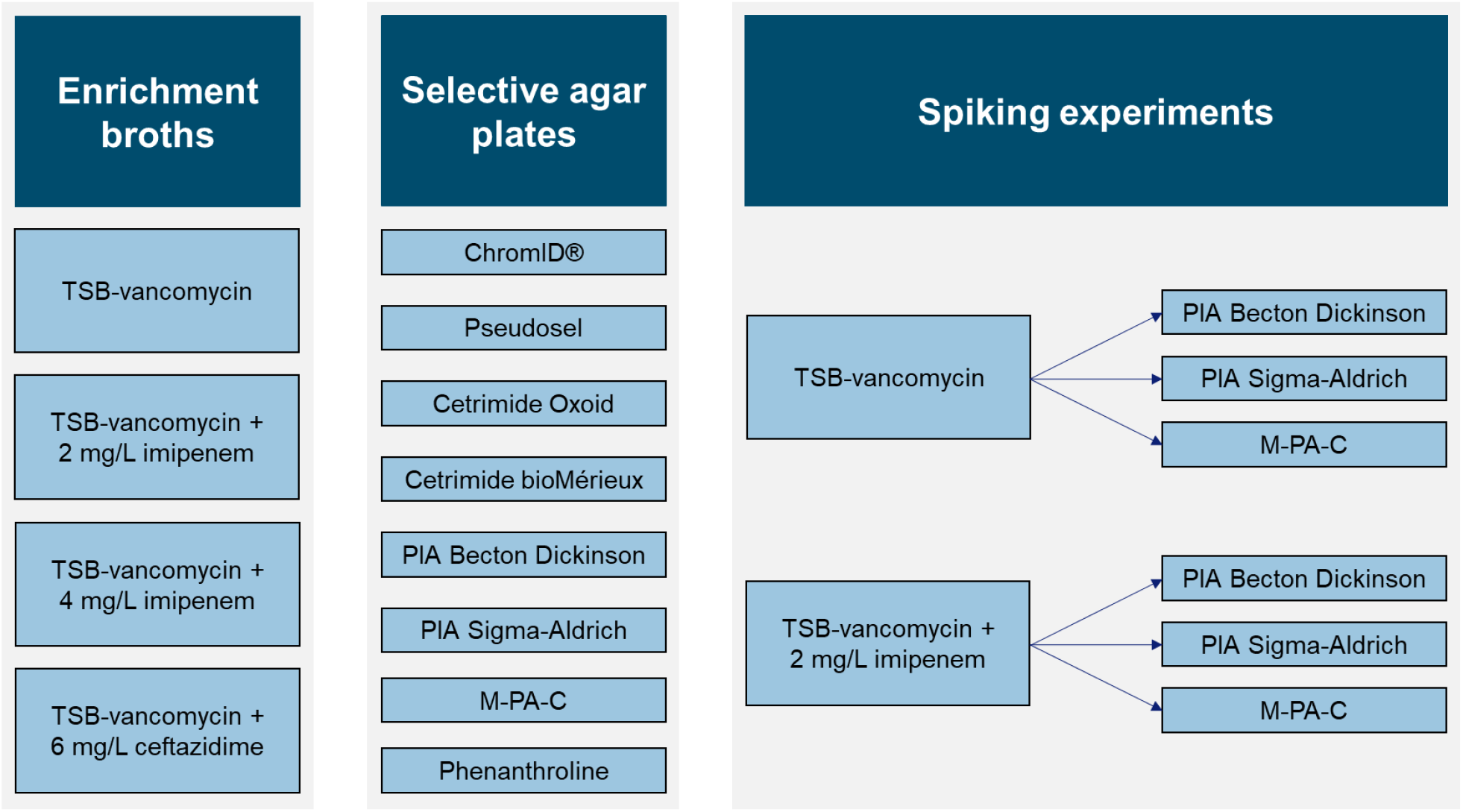
Overview of enrichment broths and agar plates tested, and spiking experiments performed. *PIA Pseudomonas* isolation agar, *TSB* tryptic soy broth.

### Comparison of selective agar plates

Eight *Pseudomonas aeruginosa*-selective culture media were tested: ChromID® *Pseudomonas aeruginosa* (bioMérieux), Pseudosel (cetrimide) agar (bioMérieux), Thermo Scientific™ *Pseudomonas* C-N Selective Agar (Oxoid, Basingstoke, UK), Cetrimide agar (bioMérieux), *Pseudomonas* isolation agar (PIA; Becton Dickinson Diagnostics, Breda, The Netherlands), PIA (Sigma-Aldrich, St. Louis, MO, USA), M-PA-C agar (Becton Dickinson Diagnostics, Breda, The Netherlands), and Phenanthroline agar (Mueller Hinton, Oxoid, Basingstoke, UK with 50 µg/L 1,10-Phenanthroline, Sigma-Aldrich, St. Louis, MO, USA) (20, 21). Blood agar plates were used as a control (Figure 1).

Bacterial suspensions (100 µL each) were inoculated directly onto each agar plate. The dilution was spread evenly over the surface of the agar plates using a disposable spreader. The agar plates were incubated aerobically at 35±1°C, followed by observation of colony characteristics after 18, 24, 42, 48, and 72 hours of incubation. Colony growth on the selective plates was recorded as growth (positive) or no growth (negative). Atypical colonies observed on the plates were identified by MALDI-TOF MS to exclude contamination. The experiment was performed in triplicate. The sensitivity was measured as the number of strains with growth divided by the number of strains tested per type of agar. The number of grown colonies was counted on each plate after 48 hours of incubation for the calculation of the yield in colonies forming unit (CFU) per mL. When the number of colonies exceeded 100, it would be scored as “> 100”.

### Detection of CRPA from spiked faeces cultured in enrichment broths with subculturing on selective agar plates

Spiking experiments were performed to further evaluate the best performing enrichment broths in combination with the best performing selective agar plates (Figure 1). Faecal samples without carbapenem-resistant bacteria were used for these experiments. Faecal solutions were prepared by suspending 5 grams of patients’ faeces into 50 mL of sterile distilled water. The spiked samples were made by adding 100 µL of the CRPA strain suspensions to 900 µL of the faecal suspension. In addition to the seven well-characterized strains of CRPA mentioned previously, suspensions of carbapenem-susceptible *P. aeruginosa* (CSPA) ATCC 27853 and *Aeromonas caviae* ATCC 15468 were also used. The latter was chosen as this microorganism is often present in water samples, and our detection method was developed in a One Health project, also focused on water. As a negative control, 100 µL of physiological salt (0.85%) was added to 900 µL of the faecal suspension.

Following the preparatory steps, 100 µL of the spiked samples and negative control were added to the selected enrichment broths and incubated for 24 hours at 35±1°C. The next day, 100 µL of the broth was sub-cultured onto the selected selective agar plates and spread evenly. The plates were then incubated for 18, 24, and 48 hours at 35±1°C. Colonies were identified using MALDI-TOF MS, followed by testing the susceptibility to carbapenems using the disc diffusion test for *P. aeruginosa* according to EUCAST. The growth of CRPA and other microorganisms on the plate was recorded. The experiment was performed three times with different faecal samples.

## RESULTS

### Comparison of enrichment broths

TSB-vancomycin and TSB-vancomycin supplemented with 2 mg/L imipenem both had 100% sensitivity with the dilutions of 10^−5^ and 10^−6^ (Table 2). Even though the sensitivity of TSB-vancomycin and TSB-vancomycin supplemented with 2 mg/L imipenem decreased to 85.7% and 71.4%, respectively, with the dilution of 10^−7^, each strain could be recovered in at least one of the experiments. For TSB-vancomycin supplemented with 4 mg/L imipenem, the sensitivity already decreased to 85.7% in the 10^−6^ dilution, whereas for the TSB-vancomcyin supplemented with 6 mg/L ceftazidime the sensitivity was only 76.2% with the 10^−5^ dilution. Overall, the sensitivities of TSB-vancomycin and TSB-vancomycin supplemented with 2 mg/L of imipenem were highest with the different dilutions and were selected for further testing.

**Table 2.** Evaluation of four different enrichment broths with seven well-characterized carbapenem-resistant *Pseudomonas aeruginosa* strains, each tested in triplo.

| Dilutions | Sensitivity |  |  |  |
| --- | --- | --- | --- | --- |
|  | TSB-vancomycin | TSB-vancomycin<br>+ 2 mg/L imipenem | TSB-vancomycin<br>+ 4 mg/L imipenem | TSB-vancomycin<br>+ 6 mg/L ceftazidime |
| $10^{-5}$ | 21/21 - 100%<br>7/7 strains | 21/21 - 100%<br>7/7 strains | 21/21 - 100%<br>7/7 strains | 16/21 - 76.2%<br>6/7 strains |
| $10^{-6}$ | 21/21 - 100%<br>7/7 strains | 21/21 - 100%<br>7/7 strains | 18/21 - 85.7%<br>7/7 strains | 13/21 - 57.1%<br>5/7 strains |
| $10^{-7}$ | 18/21 - 85.7%<br>7/7 strains | 15/21 - 71.4%<br>7/7 strains | 5/21 - 23.8%<br>4/7 strains | 7/21 - 33.3%<br>4/7 strains |
| $10^{-8}$ | 0/21 - 0%<br>0/7 strains | 3/21 - 14.3%<br>3/7 strains | 3/21 - 14.3%<br>3/7 strains | 1/21 - 4.8%<br>1/7 strains |

TSB-vancomycin tryptic soy broth supplemented with 2 mg/L of vancomycin.

### Comparison of selective agar plates

All strains were able to grow on all selective agar plates after 24 hours of incubation in experiment with the dilution of 10^−5^ (1.5 × 10^2^ CFU/mL), with no additional colonies observed on all the agar plates after 48 hours. However, *P. aeruginosa* colonies were better recognizable after 42 hours (Figure 2).

**Figure 2.**
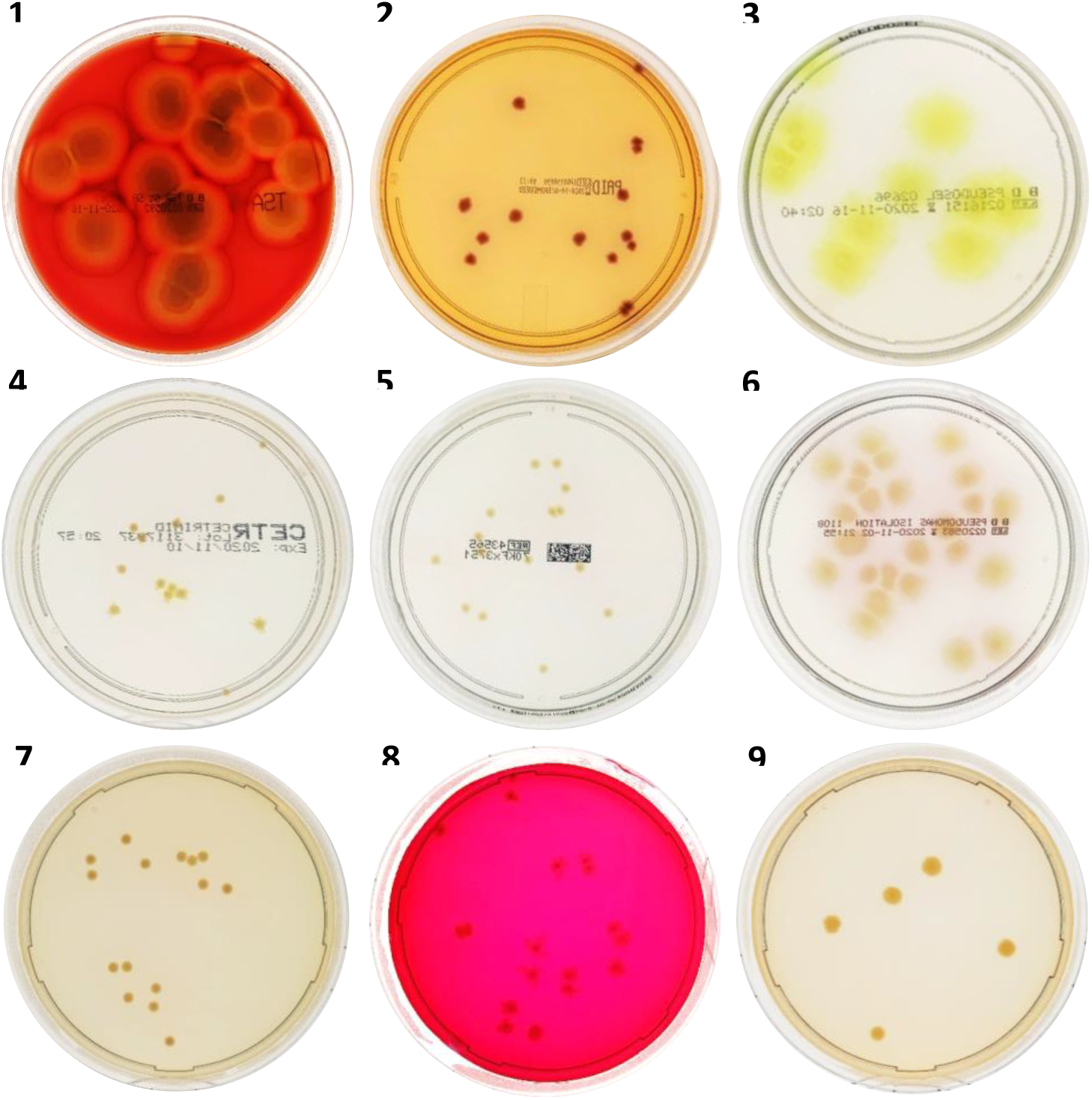
Growth of carbapenem-resistant *Pseudomonas aeruginosa* on different agar plates after 42 hours of incubation.

The agar plates shown in Figure 2 are blood agar (1), ChromID (2), Pseudosel (3), Cetrimide Oxoid (4), Cetrimide bioMérieux (5), PIA Becton Dickinson (6), PIA Sigma-Aldrich (7), M-PA-C agar (8), and Phenanthroline agar (9). The images show the growth of strain 2, 10^−6^ dilution, after 42 hours of incubation.

Figure 2 shows the colony morphologies on different selective agar plates with blood agar as the control. *P. aeruginosa* colonies were yellow-brown on Cetrimide Oxoid, Cetrimide bioMérieux, PIA Becton Dickinson, PIA Sigma-Aldrich, and Phenantroline agar plates. On ChromID agar plates, colonies had purplish-pink pigmentation with dark blue centers, while M-PA-C agar plates showed pinkish-pigmentation with dark centers. The colonies were bright greenish-yellow on the Pseudosel agar plates.

Table 3 shows the sensitivity of the selective agar plates and the blood agar plates. At 18 hours of incubation, PIA Sigma Aldrich had the highest sensitivity with the dilutions up to 10^−7^. With that dilution, PIA Sigma Aldrich was more sensitive compared to the blood agar plate (*i*.*e*., the current gold standard). After 18 hours of incubation, no additional growth was observed on the blood agar and Phenanthroline agar plates, while the sensitivity increased for the other selective agar plates. After 24 hours of incubation, there was only one additional colony growth from each cetrimide bioMérieux (observed after 42 hours of incubation), PIA Becton Dickinson and PIA Sigma-Aldrich plate (both were observed after 48 hours of incubation). No additional growth was seen for incubation periods longer than 48 hours. Compared to the blood agar plate, ChromID, Cetrimide Oxoid, PIA Becton Dickinson, PIA Sigma-Aldrich, and M-PA-C all had higher sensitivities across the different incubation times and/or dilutions. Overall, PIA Sigma Aldrich had the highest sensitivity, followed by Cetrimide Oxoid and ChromID. Pseudosel, Cetrimide bioMérieux, and Phenanthroline agar plates did not perform well and no CRPA from the 10^−8^ dilution grew. All three experiments showed consistent results.

**Table 3.** Number of strains growing on the plates.

| Dilution | ChromID | Pseudosel | Cetrimide Oxoid | Cetrimide bioMérieux | PIA BD | PIA SA | M-PA-C | Phenanthroline | Blood agar |
| --- | --- | --- | --- | --- | --- | --- | --- | --- | --- |
| **18 hours** |  |  |  |  |  |  |  |  |  |
| 10<sup>-5</sup> | 7 of 7<br>(100%) | 6 of 7<br>(86%) | 7 of 7<br>(100%) | 7 of 7<br>(100%) | 7 of 7<br>(100%) | 7 of 7<br>(100%) | 7 of 7<br>(100%) | 7 of 7<br>(100%) | 7 of 7<br>(100%) |
| 10<sup>-6</sup> | 6 of 7<br>(86%) | 6 of 7 –<br>(86%) | 6 of 7<br>(86%) | 5 of 7<br>(71%) | 7 of 7<br>(100%) | 7 of 7<br>(100%) | 7 of 7<br>(100%) | 5 of 7<br>(71%) | 7 of 7<br>(100%) |
| 10<sup>-7</sup> | 3 of 7<br>(43%) | 1 of 7 –<br>(14%) | 2 of 7<br>(29%) | No growth | 1 of 7 –<br>(14%) | 3 of 7<br>(43%) | 1 of 7<br>(14%) | 1 of 7<br>(14%) | 2 of 7<br>(29%) |
| 10<sup>-8</sup> | No growth | No growth | No growth | No growth | No growth | No growth | 1 of 7<br>(14%) | No growth | 1 of 7<br>(14%) |
| **24 hours** |  |  |  |  |  |  |  |  |  |
| 10<sup>-5</sup> | 7 of 7<br>(100%) | 7 of 7<br>(100%) | 7 of 7<br>(100%) | 7 of 7<br>(100%) | 7 of 7<br>(100%) | 7 of 7<br>(100%) | 7 of 7<br>(100%) | 7 of 7<br>(100%) | 7 of 7<br>(100%) |
| 10<sup>-6</sup> | 7 of 7<br>(100%) | 7 of 7<br>(100%) | 7 of 7<br>(100%) | 5 of 7<br>(71%) | 7 of 7<br>(100%) | 7 of 7<br>(100%) | 7 of 7<br>(100%) | 5 of 7<br>(71%) | 7 of 7<br>(100%) |
| 10<sup>-7</sup> | 4 of 7<br>(57%) | 2 of 7<br>(29%) | 4 of 7<br>(57%) | 2 of 7<br>(29%) | 2 of 7 –<br>(29%) | 5 of 7<br>(71%) | 3 of 7<br>(43%) | 1 of 7<br>(14%) | 2 of 7<br>(29%) |
| 10<sup>-8</sup> | 1 of 7<br>(14%) | No growth | 2 of 7<br>(29%) | No growth | 1 of 7 –<br>(14%) | 2 of 7<br>(29%) | No growth | No growth | 1 of 7<br>(14%) |
| **42 hours** |  |  |  |  |  |  |  |  |  |
| 10<sup>-5</sup> | 7 of 7<br>(100%) | 7 of 7<br>(100%) | 7 of 7<br>(100%) | 7 of 7<br>(100%) | 7 of 7<br>(100%) | 7 of 7<br>(100%) | 7 of 7<br>(100%) | 7 of 7<br>(100%) | 7 of 7<br>(100%) |
| 10<sup>-6</sup> | 7 of 7<br>(100%) | 7 of 7<br>(100%) | 7 of 7<br>(100%) | 5 of 7<br>(71%) | 7 of 7<br>(100%) | 7 of 7<br>(100%) | 7 of 7<br>(100%) | 5 of 7<br>(71%) | 7 of 7<br>(100%) |
| 10<sup>-7</sup> | 4 of 7<br>(57%) | 2 of 7<br>(29%) | 4 of 7<br>(57%) | 2 of 7<br>(29%) | 2 of 7 –<br>(29%) | 5 of 7<br>(71%) | 3 of 7<br>(43%) | 1 of 7<br>(14%) | 2 of 7<br>(29%) |
| 10<sup>-8</sup> | 1 of 7<br>(14%) | No growth | 2 of 7<br>(29%) | No growth | 1 of 7 –<br>(14%) | 2 of 7<br>(29%) | No growth | No growth | 1 of 7<br>(14%) |
| **48 hours** |  |  |  |  |  |  |  |  |  |
| 10<sup>-5</sup> | 7 of 7<br>(100%) | 7 of 7<br>(100%) | 7 of 7<br>(100%) | 7 of 7<br>(100%) | 7 of 7<br>(100%) | 7 of 7<br>(100%) | 7 of 7<br>(100%) | 7 of 7<br>(100%) | 7 of 7<br>(100%) |
| 10<sup>-6</sup> | 7 of 7<br>(100%) | 7 of 7<br>(100%) | 7 of 7<br>(100%) | 5 of 7<br>(71%) | 7 of 7<br>(100%) | 7 of 7<br>(100%) | 7 of 7<br>(100%) | 5 of 7<br>(71%) | 7 of 7<br>(100%) |
| 10<sup>-7</sup> | 4 of 7<br>(57%) | 2 of 7<br>(29%) | 4 of 7<br>(57%) | 2 of 7<br>(29%) | 2 of 7 –<br>(29%) | 5 of 7<br>(71%) | 3 of 7<br>(43%) | 1 of 7<br>(14%) | 2 of 7<br>(29%) |
| 10<sup>-8</sup> | 1 of 7<br>(14%) | No growth | 2 of 7<br>(29%) | No growth | 1 of 7 –<br>(14%) | 2 of 7<br>(29%) | No growth | No growth | 1 of 7<br>(14%) |
| **72 hours** |  |  |  |  |  |  |  |  |  |
| 10<sup>-5</sup> | 7 of 7<br>(100%) | 7 of 7<br>(100%) | 7 of 7<br>(100%) | 7 of 7<br>(100%) | 7 of 7<br>(100%) | 7 of 7<br>(100%) | 7 of 7<br>(100%) | 7 of 7<br>(100%) | 7 of 7<br>(100%) |
| 10<sup>-6</sup> | 7 of 7<br>(100%) | 7 of 7<br>(100%) | 7 of 7<br>(100%) | 5 of 7<br>(71%) | 7 of 7<br>(100%) | 7 of 7<br>(100%) | 7 of 7<br>(100%) | 5 of 7<br>(71%) | 7 of 7<br>(100%) |
| 10<sup>-7</sup> | 4 of 7<br>(57%) | 2 of 7<br>(29%) | 4 of 7<br>(57%) | 2 of 7<br>(29%) | 2 of 7 –<br>(29%) | 5 of 7<br>(71%) | 3 of 7<br>(43%) | 1 of 7<br>(14%) | 2 of 7<br>(29%) |
| 10<sup>-8</sup> | 1 of 7<br>(14%) | No growth | 2 of 7<br>(29%) | No growth | 1 of 7 –<br>(14%) | 2 of 7<br>(29%) | No growth | No growth | 1 of 7<br>(14%) |

Table 4 shows the number of CFU/mL for each strain on each agar plate. For some strains, some agar plates failed to yield a higher number of colonies than the standard blood agar. For instance, phenanthroline agar only yielded a higher number of colonies of the strain 1. None of the selective agar plates yielded a higher number of colonies of strain 7 compared to blood agar. On average, the M-PA-C agar had the highest yield (1.53), while the Phenanthroline agar had the lowest yield (0.55). There were three agar plates with higher yields than blood agar, M-PA-C, PIA Becton Dickinson and PIA Sigma-Aldrich. Based on the combination of sensitivity and yield of each plate, those three were selected for the spiking experiments.

**Table 4.** Yields of growth on plates after 48 hours of incubation.

| Strain | Yield (x $10^{-6}$ CFU/mL) | | | | | | | | |
| --- | --- | --- | --- | --- | --- | --- | --- | --- | --- |
|  | Blood Agar | ChromID | Pseudosel | Cetrimide Oxoid | Cetrimide bioMérieux | PIA BD | PIA SA | M-PA-C | Phenanthroline |
| 1 | 1.10 | 1.10 | 1.23 | 1.60 | 1.07 | 1.40 | 1.17 | 1.60 | 1.60 |
| 2 | 1.77 | 0.87 | 0.87 | 0.95 | 1.28 | 1.97 | 1.37 | 1.93 | 0.50 |
| 3 | 1.23 | 1.07 | 1.57 | 0.93 | 1.40 | 1.67 | 1.27 | 1.33 | 0.78 |
| 4 | 0.60 | 0.74 | 0.06 | 0.63 | 0.30 | 0.98 | 0.94 | 0.92 | 0.04 |
| 5 | 1.50 | 1.80 | 1.95 | 2.10 | 1.60 | 1.80 | 2.15 | 2.10 | 0.29 |
| 6 | 2.10 | 1.00 | 0.91 | 1.00 | 1.90 | 1.50 | 1.75 | 2.30 | 0.61 |
| 7 | 1.38 | 0.65 | 0.40 | 0.65 | 0.49 | 0.93 | 1.10 | 0.55 | 0.01 |
| Average | 1.38 | 1.03 | 1.00 | 1.12 | 1.15 | 1.46 | 1.39 | 1.53 | 0.55 |

*BD* Becton Dickinson, *PIA Pseudomonas* Isolation Agar, *SA* Sigma Aldrich.
Green = higher yield than blood agar (control)

### Growth of CRPA from human faecal samples spiked with CRPA (spiking experiment)

For these experiments, the two most sensitive enrichments broths (*i*.*e*., TSB-vancomycin and TSB-vancomycin with 2 mg/L imipenem) and the most sensitive selective agar plates (*i*.*e*., PIA Becton Dickinson, PIA Sigma-Aldrich, and M-PA-C) were combined (Table 5). All selective agar plates used in combination with TSB-vancomycin with 2 mg/L imipenem had the same sensitivity in all samples. The combination of TSB-vancomycin and M-PA-C had the highest sensitivity in detecting CRPA in Sample 1 and 3. No CSPA was detected when TSB-vancomycin with 2 mg/L imipenem was used. As expected, CSPA was found when using a broth without antibiotics, which hampered growth of CRPA. In all samples and all experiments, there was no *Pseudomonas* spp. other than *P. aeruginosa* found. The agar plate with the least amount of other growth was M-PA-C. All six methods allowed yeasts, such as *Candida albicans* and *Candida tropicalis*, to grow.

**Table 5.**
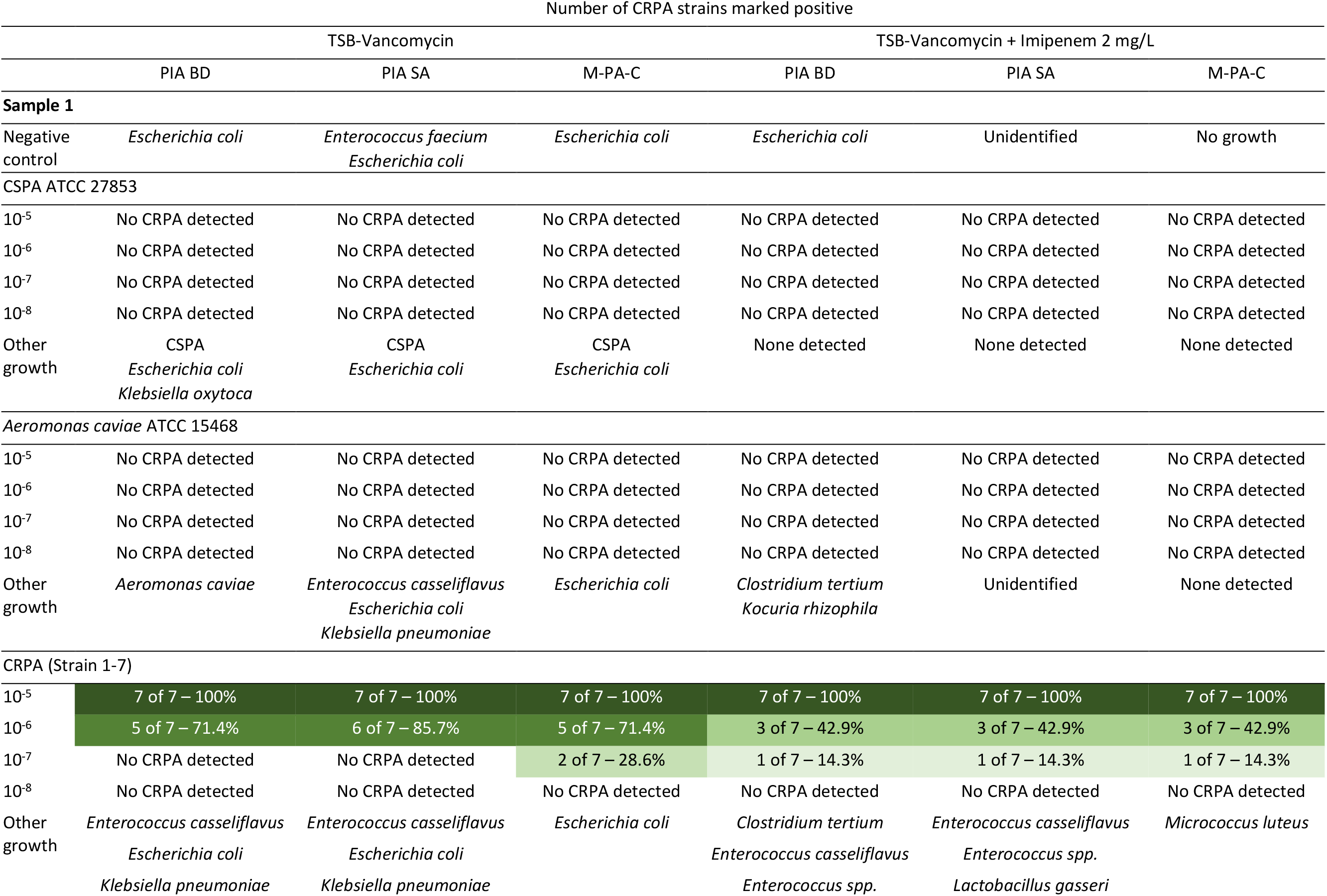
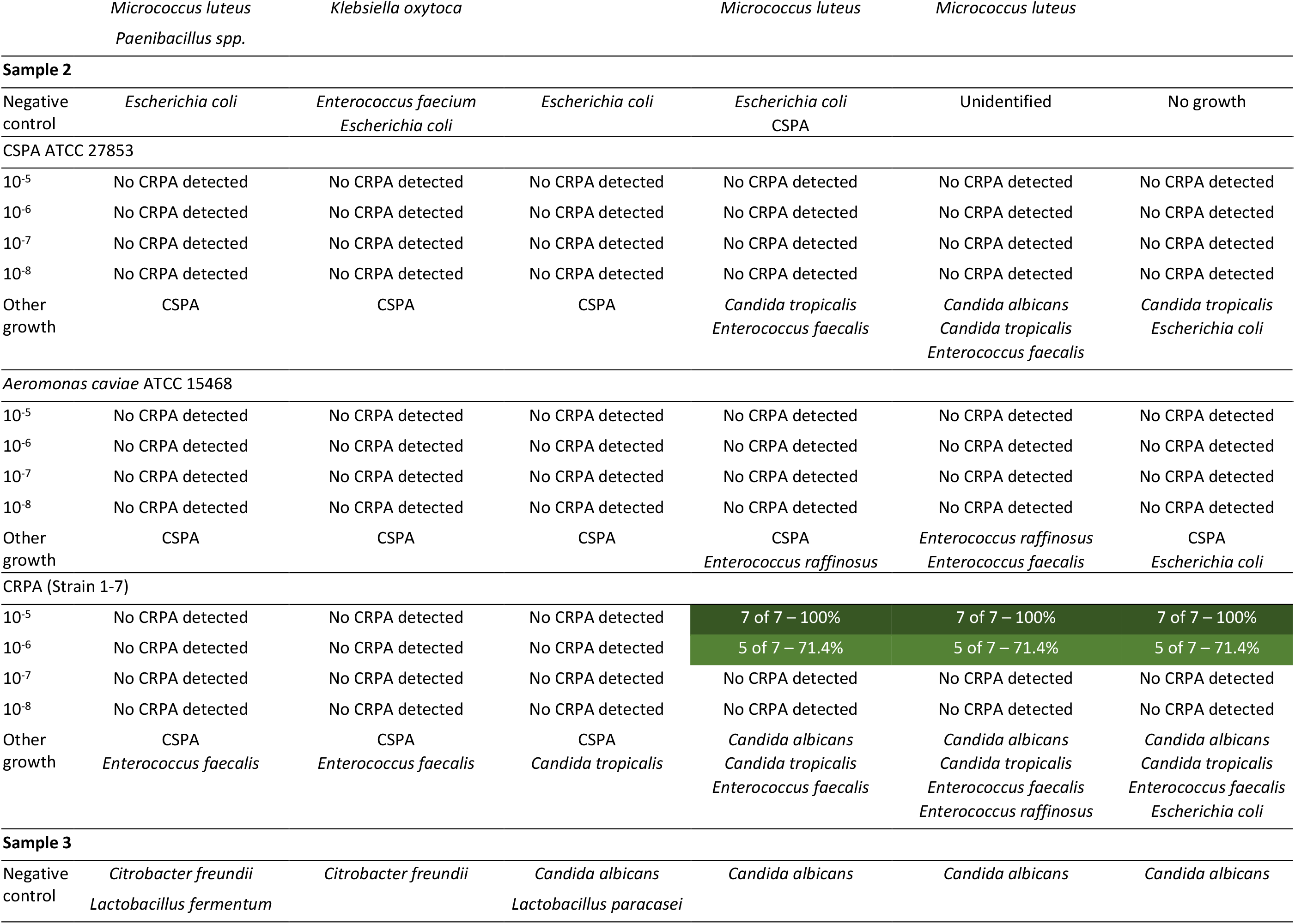
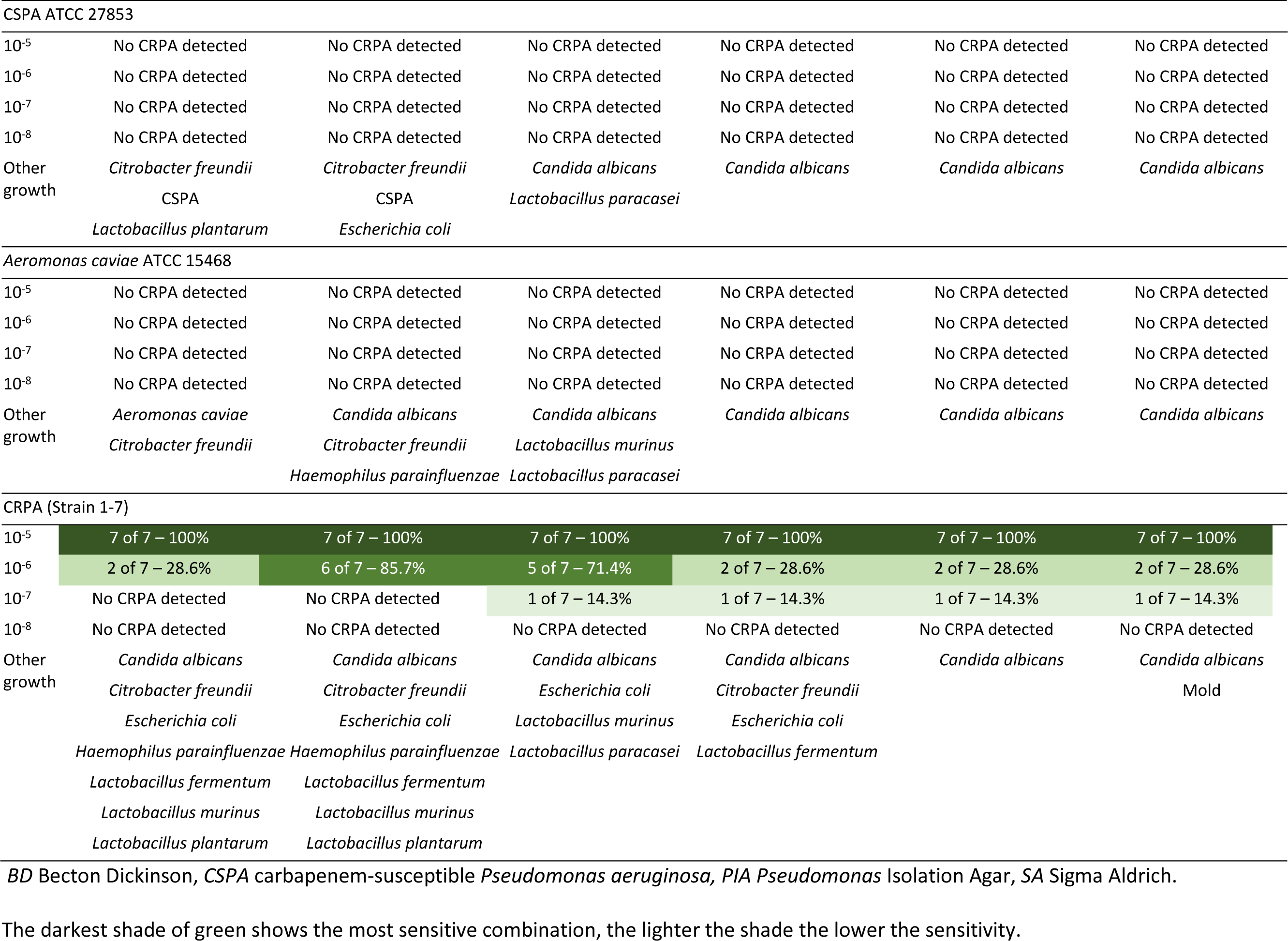
Evaluation of the combination of different enrichment broths and selective agar plates for the detection of carbapenem-resistant *Pseudomonas aeruginosa*.

## DISCUSSION

This study shows that the best method for the detection of CRPA is by inoculating the sample in TSB -vancomycin supplemented with 2 mg/L imipenem, continued by subculturing the broth onto the M-PA-C agar plate. Imipenem supplementation in the enrichment broth was efficiently eliminating CSPA, even when CSPA was intentionally added. When the sample naturally contained CSPA, it masked the CRPA spiked into the sample. As a result, the methods without imipenem supplementation failed to grow the CRPA. Despite having a lower sensitivity in higher dilutions, the combination of TSB-vancomycin with imipenem and M-PA-C agar plate resulted in the least amount of growth of other microorganisms.

Use of a selective enrichment broth has proven to be useful for the detection of carriage of various multidrug-resistant microorganisms, such methicillin-resistant *Staphylococcus aureus* and vancomycin-resistant *Enterococcus faecium* (22, 23). For CRPA, there is only one report evaluating the use of an enrichment broth supplemented with antibiotics (*i*.*e*., meropenem) (24). However, as mentioned in the methods section, imipenem was chosen as the carbapenem of choice, because a study reported its stability compared to meropenem (19).

Selective agar plates are useful as they promote growth of a specific microorganism, in our case *P. aeruginosa*, while inhibiting other species. Thus, identification of CRPA would be feasible. Moreover, the increased pigment production resulting from culturing *P. aeruginosa* on selective agar plates containing magnesium chloride and potassium sulfate makes the identification easier (Supplementary Table 1). Among eight different selective agar plates tested, M-PA-C and PIA Sigma-Aldrich showed high sensitivities (Table 3) and yields (1.53 and 1.39 × 10^−6^ CFU/mL, respectively). One of the differences between these two media is the addition of nalidixic acid to inhibit other Gram-negative bacteria and kanamycin as selective agent to inhibit the growth of Gram-positive bacteria in the M-PA-C agar. A previous study showed that medium containing nalidixic acid and kanamycin has a higher sensitivity in detecting *P. aeruginosa* compared to other selective media (21).

In the spiking experiment using TSB-vancomycin, the growth of *E. coli* was found in sample 1 and sample 3 on all agar plates tested. *E. coli* is generally susceptible to nalidixic acid (selective component of M-PA-C) and some strains of *E. coli* are reported to be resistant against triclosan (selective component of PIA) (25, 26). In the manufacturers’ guide of all agar plates tested, *E. coli* ATCC 25922 is reported as the quality control strain and should result in partial to complete inhibition. When imipenem was added to the enrichment broth, the growth of *E. coli* was inhibited and only detected in sample 2 (on M-PA-C) and sample 3 (on PIA Becton Dickinson plate). Yeast could not be eliminated by the six methods tested and could potentially interfere with CRPA growth. Colonies, however, can be easily recognized as yeasts.

Spiking experiments simulate how the method performs when applied to human samples. Numerous bacteria in the normal flora may obscure the low number of CRPA in nonselective culture methods. When adding either CSPA or *A. caviae*, it was shown that these were suppressed by the methods with TSB-vancomycin supplemented with imipenem 2 mg/L.

The broth used in this study can be made by adding antibiotic discs to the broth, generally used in laboratories around the world. The selective agar with the highest sensitivity in this study can be shipped and stored easily because it is sold as powder. Thus, the proposed method is feasible for laboratories with limited resources or in remote areas. Furthermore, because of its high sensitivity, this method can be used in surveillance or screening of CRPA in healthy people as well.

This study has some limitations. First, *Pseudomonas* spp. other than *P. aeruginosa* were not included in the spiking experiment. The selective agar plates used in this study are selective for *P. aeruginosa* and supposedly able to suppress the growth of those species or at least make the other colonies colorless. Second, only seven strains have been used in this study, but they all are important high-risk clones of CRPA. Third, our study was focused on screening with rectal or faecal samples and we did not include other body sites which can be useful for screening, such as throat. Finally, faecal samples used in the spiking experiment were from Dutch persons and their microbiota might be different from persons in other countries.

## CONCLUSION

In this study, a highly-sensitive method to detect carriage of CRPA was developed using a stepwise approach. The best method for detecting CRPA based on sensitivity and selectivity was the combination of TSB-vancomycin with 2 mg/L imipenem as an enrichment broth for overnight incubation, followed by subculturing the broth onto an M-PA-C agar plate. Careful colony selection followed by identification and susceptibility testing is needed after a positive result to confirm the screening results. Real implementation of the screening of CRPA in humans, where the number of CRPA might be limited and affected by a variety of normal flora, is needed to verify the clinical use and the practicality.

## CRediT AUTHORSHIP CONTRIBUTION STATEMENT

S.N.S.: formal analysis, visualization, writing – original draft, writing – review & editing

A.V.: project administration, writing – review & editing

N.K: investigation, resources, validation, writing – review & editing

A.R.: investigation, resources, validation, writing – review & editing

H.S.: supervision, writing – review & editing

Y.R.S: writing – review & editing

M.C.V.: supervision, writing – review & editing

A.K.: funding acquisition, supervision, writing – review & editing

J.A.S.: conceptualization, formal analysis, funding acquisition, methodology, supervision, validation, writing – review & editing

## ACKNOWLEDGEMENTS

The authors are grateful to all members of the SAMPAN Consortium for their input: Anniek de Jong (Deltares, Delft, the Netherlands) and Roger C. Lévesque (U. Laval Integrative Systems Biology Institute, Québec, Canada). Also, the authors would like to acknowledge the National Institute for Infectious Diseases “L. Spallanzani” IRCCS, Rome, Italy, for their contribution in the study design (Enrico Girardi).

## REFERENCES

1. Qin S, Xiao W, Zhou C, Pu Q, Deng X, Lan L, et al. Pseudomonas aeruginosa: pathogenesis, virulence factors, antibiotic resistance, interaction with host, technology advances and emerging therapeutics. Signal Transduct Target Ther. 2022;7(1):199.

2. Saharman YR, Pelegrin AC, Karuniawati A, Sedono R, Aditianingsih D, Goessens WHF, et al. Epidemiology and characterisation of carbapenem-non-susceptible Pseudomonas aeruginosa in a large intensive care unit in Jakarta, Indonesia. Int J Antimicrob Agents. 2019;54(5):655–60.

3. Persoon MC, Voor In ‘t Holt AF, van Meer MPA, Bokhoven KC, Gommers D, Vos MC, et al. Mortality related to Verona Integron-encoded Metallo-β-lactamase-positive Pseudomonas aeruginosa: assessment by a novel clinical tool. Antimicrob Resist Infect Control. 2019;8:107.

4. WHO Bacterial Priority Pathogens List, 2024: bacterial pathogens of public health importance to guide research, development and strategies to prevent and control antimicrobial resistance. Geneva: World Health Organization; 2024.

5. Pettigrew MM, Gent JF, Kong Y, Halpin AL, Pineles L, Harris AD, et al. Gastrointestinal microbiota disruption and risk of colonization with carbapenem-resistant Pseudomonas aeruginosa in intensive care unit patients. Clin Infect Dis. 2019;69(4):604–13.

6. Pirzadian J, Harteveld SP, Ramdutt SN, van Wamel WJB, Klaassen CHW, Vos MC, et al. Novel use of culturomics to identify the microbiota in hospital sink drains with and without persistent VIM-positive Pseudomonas aeruginosa. Sci Rep. 2020;10(1):17052.

7. Büchler AC, Shahab SN, Severin JA, Vos MC, Voor In ‘t Holt AF. Outbreak investigations after identifying carbapenem-resistant Pseudomonas aeruginosa: a systematic review. Antimicrob Resist Infect Control. 2023;12(1):28.

8. Gbaguidi-Haore H, Varin A, Cholley P, Thouverez M, Hocquet D, Bertrand X. A Bundle of Measures to Control an Outbreak of Pseudomonas aeruginosa Associated with P-Trap Contamination. Infect Control Hosp Epidemiol. 2018;39(2):164–9.

9. Slekovec C, Robert J, van der Mee-Marquet N, Berthelot P, Rogues AM, Derouin V, et al. Molecular epidemiology of Pseudomonas aeruginosa isolated from infected ICU patients: a French multicenter 2012–2013 study. Eur J Clin Microbiol Infect Dis. 2019;38(5):921–6.

10. Shahab SN, van Veen A, Büchler AC, Saharman YR, Karuniawati A, Vos MC, et al. In search of the best method to detect carriage of carbapenem-resistant Pseudomonas aeruginosa in humans: a systematic review. Ann Clin Microbiol Antimicrob. 2024;23(1):50.

11. Carbapenem-resistant Pseudomonas Aeruginosa: the SAMPAN study. https://classic.clinicaltrials.gov/show/NCT05282082.

12. Matuschek E, Brown DF, Kahlmeter G. Development of the EUCAST disk diffusion antimicrobial susceptibility testing method and its implementation in routine microbiology laboratories. Clin Microbiol Infect. 2014;20(4):O255–66.

13. EUCAST. Breakpoint tables for interpretation of MICs and zone diameters, version 8.1, 2018 [Available from: http://www.eucast.org/ast_of_bacteria/previous_versions_of_documents/.

14. EUCAST. Breakpoint tables for interpretation of MICs and zone diameters, version 10.0, 2020 [updated 2020. Available from: http://www.eucast.org/ast_of_bacteria/previous_versions_of_documents/.

15. van der Zee A, Kraak WB, Burggraaf A, Goessens WHF, Pirovano W, Ossewaarde JM, et al. Spread of carbapenem resistance by transposition and conjugation among Pseudomonas aeruginosa. Front Microbiol. 2018;9:2057.

16. Pirzadian J, Persoon MC, Severin JA, Klaassen CHW, de Greeff SC, Mennen MG, et al. National surveillance pilot study unveils a multicenter, clonal outbreak of VIM-2-producing Pseudomonas aeruginosa ST111 in the Netherlands between 2015 and 2017. Sci Rep. 2021;11(1):21015.

17. Pirzadian J, Voor In ‘t Holt AF, Hossain M, Klaassen CHW, de Goeij I, Koene H, et al. Limiting spread of VIM-positive Pseudomonas aeruginosa from colonized sink drains in a tertiary care hospital: A before-and-after study. PLoS One. 2023;18(3):e0282090.

18. Pelegrin AC, Saharman YR, Griffon A, Palmieri M, Mirande C, Karuniawati A, et al. High-risk international clones of carbapenem-nonsusceptible Pseudomonas aeruginosa endemic to Indonesian intensive care units: impact of a multifaceted infection control intervention analyzed at the genomic level. mBio. 2019;10(6).

19. Shahab SN, Bexkens ML, Kempenaars N, Rijfkogel A, Karuniawati A, Vos MC, et al. Stability of four carbapenem antibiotics in discs used for antimicrobial susceptibility testing. bioRxiv. 2024:2024.06.17.599257.

20. Keeven JK, DeCicco BT. Selective medium for Pseudomonas aeruginosa that uses 1,10-phenanthroline as the selective agent. Appl Environ Microbiol. 1989;55(12):3231–3.

21. Kodaka H, Iwata M, Yumoto S, Kashitani F. Evaluation of a new agar medium containing cetrimide, kanamycin and nalidixic acid for isolation and enhancement of pigment production of Pseudomonas aeruginosa in clinical samples. J Basic Microbiol. 2003;43(5):407–13.

22. Safdar N, Narans L, Gordon B, Maki DG. Comparison of culture screening methods for detection of nasal carriage of methicillin-resistant Staphylococcus aureus: a prospective study comparing 32 methods. J Clin Microbiol. 2003;41(7):3163–6.

23. Murk JL, Heddema ER, Hess DL, Bogaards JA, Vandenbroucke-Grauls CM, Debets-Ossenkopp YJ. Enrichment broth improved detection of extended-spectrum-beta-lactamase-producing bacteria in throat and rectal surveillance cultures of samples from patients in intensive care units. J Clin Microbiol. 2009;47(6):1885–7.

24. Fournier C, Poirel L, Nordmann P. Implementation and evaluation of methods for the optimal detection of carbapenem-resistant and colistin-resistant Pseudomonas aeruginosa and Acinetobacter baumannii from stools. Diagn Microbiol Infect Dis. 2020;98(2):115121.

25. Zechner V, Sofka D, Paulsen P, Hilbert F. Antimicrobial resistance in Escherichia coli and resistance genes in coliphages from a small animal clinic and in a patient dog with chronic urinary tract infection. Antibiotics (Basel). 2020;9(10).

26. Yu BJ, Kim JA, Pan JG. Signature gene expression profile of triclosan-resistant Escherichia coli. J Antimicrob Chemother. 2010;65(6):1171–7.

