## Supplementary table 1 for "Development of a highly-sensitive method to detect the carriage of carbapenem-resistant *Pseudomonas aeruginosa* in humans"

| Composition | ChromID | Pseudosel | Cetrimide Oxoid | Cetrimide bioMérieux* | PIA BD | PIA SA | M-PA-C | Phenan-throline |
|---|---|---|---|---|---|---|---|---|
| Macronutrients | | | | | | | | |
| Casein peptone (bovine) | 13.0 g | | | | | | | |
| Meat peptone (porcine) | 6.0 g | | | | | | | |
| Pancreatic digest of gelatin (gelatin peptone) | | 20.0 g | 16.0 g | | | | | |
| Protein digest of casein (casein hydrolysate) | | | 10.0 g | | | | | |
| Bacto peptone | | | | | 20.0 g | 20.0 g | | |
| Beef infusion | | | | | | | | 300 g |
| Acid hydrolysate of casein | | | | | | | | 17.5 g |
| Starch | | | | | | | | 1.5 g |
| L-lysine HCl | | | | | | | 5.0 g | |
| Yeast extract | 3.5 g | | | | | | 2.0 g | |
| Lactose | | | | | | | 1.25 g | |
| Sucrose | | | | | | | 1.25 g | |
| Xylose | | | | | | | 1.25 g | |
| Amino acids | 1.0 g | | | | | | | |
| Selective agents | | | | | | | | |
| Antibiotic mixture | 0.023 g | | | | | | | |
| Susbtrate mixture | 0.125 g | | | | | | | |
| Magnesium chloride | | 1.4 g | 1.4 g | | 1.4 g | 1.4 g | | |
| Magnesium sulfate | | | | | | | 1.5 g | |
| Potassium sulfate | | 10.0 g | 10.0 g | | 10.0 g | 10.0 g | | |
| Sodium chloride | | | | | | | 5.0 g | |
| Sodium thiosulfate | | | | | | | 5.0 g | |
| Cetrimide | | 0.3 g | 0.2 g | | | | | |
| Ferric ammonium citrate | | | | | | | 0.8 g | |
| Triclosan (Irgasan®) | | | | | 0.025 g | 0.025 g | | |
| Nalidixic acid | | | | | | | 0.037 g | |

| | | | | | | | | |
|---|---|---|---|---|---|---|---|---|
| Sodium nalidixate | | | 0.015 g | | | | | |
| Kanamycin | | | | | | | 8.0 mg | |
| Phenol red | | | | | | | 0.08 g | |
| 1-10 Phenanthroline | | | | | | | | 50 µg |
| General compositions | | | | | | | | |
| Agar | 15.0 g | 13.6 g | 11.5 g | | 13.6 g | 13.6 g | 12.0 g | 17.0 g |
| Glycerol | | 10.0 mL | 10.0 mL | | 20.0 mL | | | |
| Purified water | 1 L | 1 L | 1 L | | 1 L | 1 L | 1 L | 1 L |
| pH | 6.8 | | 7.1 | | 7.0 | 7.0 | 7.2 | |
| Package | Poured plate | Poured plate | Poured plate | Poured plate | Poured plate | Powder | Powder | Powder with supplement |
| Expected *Pseudomonas aeruginosa* colony color | Pink to violet | Blue-green to green | Blue-green fluorescent or red-brown | Pale green or green fluorescent | Green to blue-green, diffuses into medium | Green or blue to blue-green | Pink or pinkish brown | |

*information is not provided
